# Label-free detection of individual virus-infected cells using deep learning

**DOI:** 10.64898/2026.01.15.699499

**Authors:** Juliane Pfeil, Corinna Siegmund, Eva Müller, Shakhnaz Akhmedova, Alexandra Löwe, Anne Kauter, Tobias Tertel, Bernd Giebel, Michael Laue, Vu Thuy Khanh Le-Trilling, Christian Sieben, Mirko Trilling, Roland Schwarzer, Nils Körber

## Abstract

Numerous applications in research and medicine rely on reliable identification and quantification of virus-infected cells. Current methods either apply reporter viruses, that often differ from clinical isolates (e. g. cell tropism, immune evasion) or staining approaches, that prevent live-cell experiments and may introduce biases through manual counting. We present a deep learning model for the label-free identification of virus-infected cells on light microscopy images (VAIruScope). To overcome limitations, our pipeline enables an automated quantification of virus-infected cells based on the recognition of cytopathic effects. The method was applied to different cell models and four clinically relevant prototype viruses representing RNA- (influenza A virus), DNA- (human cytomegalovirus, herpes simplex virus-1) and retroviruses (human immunodeficiency virus-1). VAIruScope identified infected cells achieving classification accuracies of up to 96 %. As proof-of-concept, the method was validated using electron microscopy for a wild-type HSV-1. VAIruScope may be applicable to live-cell imaging to investigate infection dynamics.

## Introduction

As obligate intracellular pathogens, viruses massively reprogram their host cells during replication [1–4]. Although the specific remodeling of infected cells varies between viruses, it is usually accompanied by dramatic subcellular and cell-morphological changes [5–7] such as for example the formation of specific replication organelles [1, 8]. These so-called virus-induced cytopathic effects (CPE) can be observed microscopically by trained personal constituting the basis of traditional virological methods such as the determination of plaque-forming units (PFU) or the tissue culture infectious dose 50 (TCID50). Moreover, individual infected cells can be identified by electron microscopy or by expensive and labor-intensive staining procedures recognizing viral nucleic acids or virus-encoded proteins (e. g., using enzyme-coupled antibodies and enzymatic reactions)[9]. Alternatively, genetically-modified viruses encoding fluorescent reporter genes such as enhanced green fluorescent protein (eGFP) can be applied [10–12]. Despite their broad application and undisputable relevance in virus research, aforementioned methods have important disadvantages: they are time-consuming, prone to errors and biases in case that operator’s evaluations are involved, and rely either on the usage of genetically-modified viruses, which do not exist for emerging viruses and often do not recapitulate all characteristics of clinical virus isolates or rely on end-point determinations ruling out live-cell imaging studies addressing infection dynamics. Importantly, single-cell CPE are highly pathogen- and host-dependent and can manifest as diverse ultrastructural alterations and their visibility is influenced by the time after infection. This heterogenous appearance complicates localization by standard brightfield (BF) microscopy, as the cell bodies and nuclei are highly divers, morphologically complex and exhibit low contrast. Consequently, human examiners are often only able to identify an obvious CPE at the single-cell level early after infection, if it is very pronounced and appears prototypic, and subtle virus-specific alterations may be missed entirely at light-microscopic resolution.

Advanced image recognition techniques based on deep learning (DL) are now broadly applied to analyze microscopic images. These methods have been instrumental in the automation and acceleration of complex tasks such as cell classification, counting, tracking, cell volume measurements, vessel segmentation and others [13]. In the field of virology, neural networks have been employed for a variety of applications, including the segmentation of infected nuclei [14, 15], the tracking of fluorescence-labelled viral particles [16], image denoising [17], and the classification of infected cells based on DNA staining in the cell nucleus [18]. Moreover, specialized microscopic techniques can be applied for the study of unlabeled virus infections, and various approaches (e. g. transmitted light microscopy, atomic force microscopy) have been used for this task [19].

Most of the research has focused on the inspection of images produced by high-throughput microscopy of multiple cells in microtiter plates. For instance, DL-based image classification models have been used to recognize an infection in cell culture across a range of viruses without specifically identifying individual infected cells [20–24]. Alternative methods have focused on changes in cell confluence or morphology, achieving high accuracy but often requiring expensive, specialized equipment and proprietary software [25, 26]. In contrast, there has been limited research focusing on the detection of infections at the single-cell level. One method used digital holo-tomography microscopy to detect CPE in single cells, but it was limited by the need for continuous recording and significant instrumental effort [27]. Another approach used the fluorescence signal with DL to identify morphological changes of the cell nuclei [18]. Although effective, this method requires manual labeling and fluorescent staining, which excludes the approach for several applications.

Recent developments in the field have therefore focused predominantly on identifying population-level CPE using BF images or on the identification of individual infected cells using specific fluorophores. Here, we present a framework that combines the advantages of both approaches for the detection of virus-infected cells with artificial intelligence (AI) and light microscopy (VAIruScope). Our model identifies individual, virus-infected cells from BF/phase contrast (PC) microscopic images without fluorescence markers or other labeling. The algorithm is able to detect CPE very early after infection (~ from day one) and accurately represents the viral dose and the effect of anti-as well as pro-viral compounds.

## Results

### Deep learning for the identification of infection status and recognition of cell centers

In order to automatically locate and classify cells according to their infection status, we developed a transformer-convolutional-hybrid neural network (called VAIruScope) that precisely predicts the cell infection status and corresponding cell center location. For model training, which utilized 80 % of the available data, we used different cell lines that had either been infected with eGFP- or yellow fluorescent protein (YFP)-labelled reporter viruses or left uninfected (mock treatment). Respective cells were fixed and for three virus/cell combinations the nuclear DNA was stained with DAPI. Different image channels were captured: the BF/PC as model input and fluorescence channels for the viral fluorescent reporter protein (eGFP/YFP) and optional nuclear staining (DAPI) as depicted in Figure 1A. VAIruScope comprises an image encoder and two parallel streams: one to determine the infection signal and one to predict the cell centers. The combination of predictions from both streams enables the assessment of the infection status and the localization of individual cells. The BF/PC channel was used as the model input, as well as the infection status based on the signal of the fluorescent protein and the center locations based on the DAPI or BF channel as the labels during training (Figure 1B). The trained VAIruScope model was capable of detecting infected cells without the need for eGFP/YFP signals or DAPI staining, exclusively using the BF/PC channel of a given cell culture experiment as input, with its performance validated on a separate, previously unseen dataset (Figure 1C).

**Figure 1.**
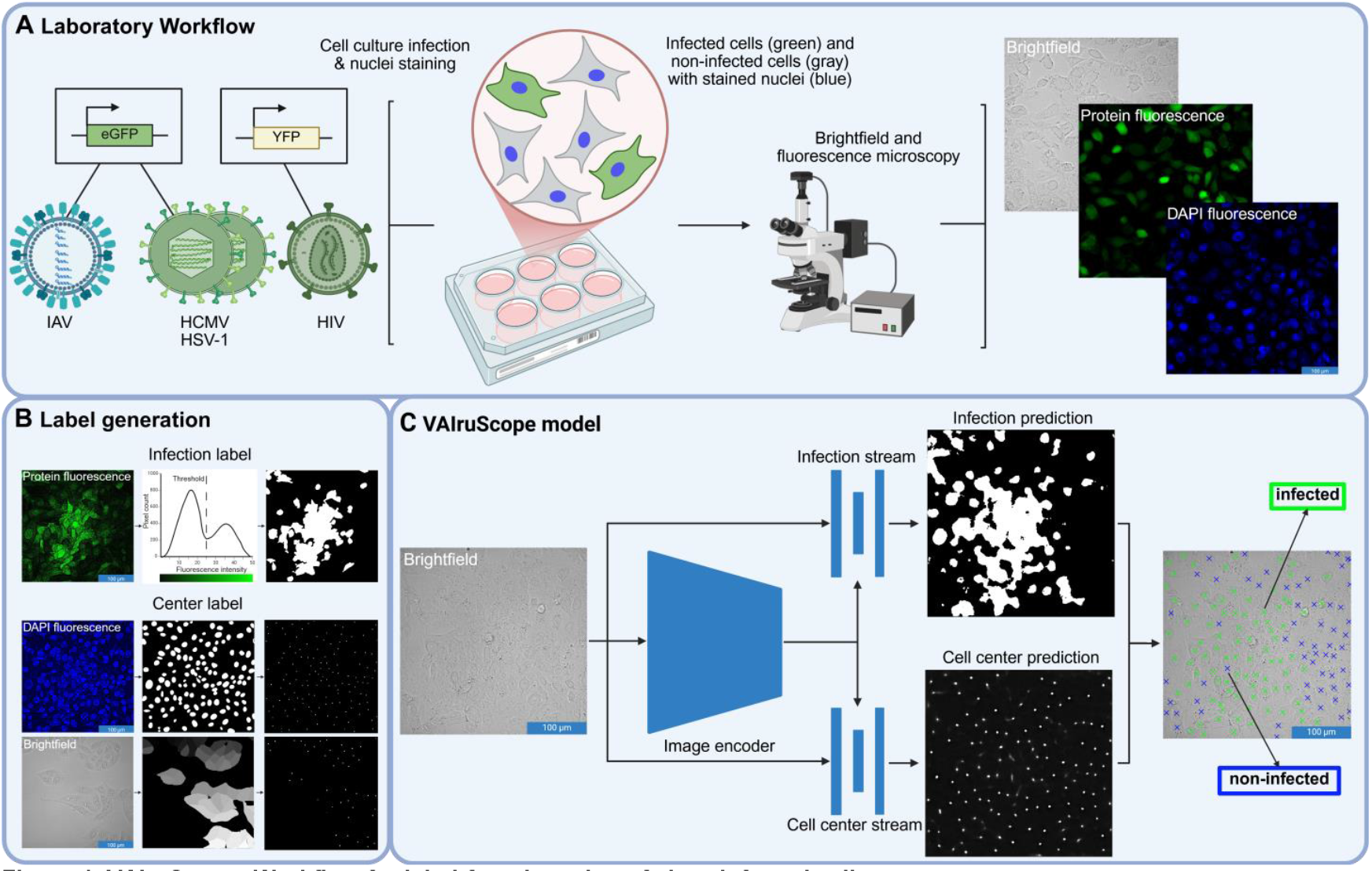
VAIruScope: Workflow for label-free detection of virus-infected cells. (A) Cultured cells were separately infected with four types of eGFP- or YFP-labelled reporter viruses (influenza A virus (IAV), human cytomegalovirus (HCMV), herpes simplex virus 1 (HSV-1) and human immunodeficiency virus (HIV-1)). Nuclei of respective cells were optionally stained with DAPI. BF/PC and fluorescence microscopy recorded multi-channel images containing BF/PC, eGFP/YFP and, where applicable, DAPI signals. (B) Infection labels were generated based on the eGFP/YFP images to generate binary infection masks. Center labels were generated based on DAPI or BF images. Individual nuclei/cells were detected and peaks with center localizations were generated. (C) VAIruScope is based on an image encoder that is fed with BF/PC images and two parallel prediction streams. The first stream utilized BF/PC images and infection labels to predict the fluorescence signal. The second stream of the DL model was trained with BF/PC images and center labels to predict center locations. The results of both streams are combined to generate a prediction for the infection status and cell location based on the BF/PC input image.

### VAIruScope reliably distinguishes between non-infected cells and cells infected with RNA-, DNA- or retroviruses

Microscopic BF/PC and fluorescence images (either at 20x or 40x magnification) were acquired to assess infections with four clinically-relevant prototype viruses representing RNA-, DNA- or retroviruses. We investigated influenza A virus (IAV), human cytomegalovirus (HCMV), herpes simplex virus 1 (HSV-1) as well as human immunodeficiency virus (HIV-1) and different cell models (A549 for IAV, MRC-5 for HCMV, BJ-5ta and Vero for HSV-1, TZM-bl for HIV-1). Cells were infected with various virus doses (low, middle, high: see details in the method section) or left uninfected (mock) and inspected at different time points post infection (1-8 days post infection (dpi)). In addition, Interferon-γ (IFN-γ) and Ruxolitinib (Ruxo) were applied to interrogate the influence of known anti-viral and pro-viral effects, respectively. By fundamentally changing the proteome of conditioned cells [28], IFN-γ elicits direct anti-viral properties especially against DNA viruses [29] including HSV-1 [30] and HCMV [31], while Ruxolitinib is a potent Janus kinase (JAK) inhibitor [32] that prevents IFN-induced signaling and its down-stream induction of anti-viral innate immunity [32] including the antiviral activity of IFN against cytomegaloviruses [33]. The dimensions and characteristics of the generated datasets are outlined in Table 1.

**Table 1.**
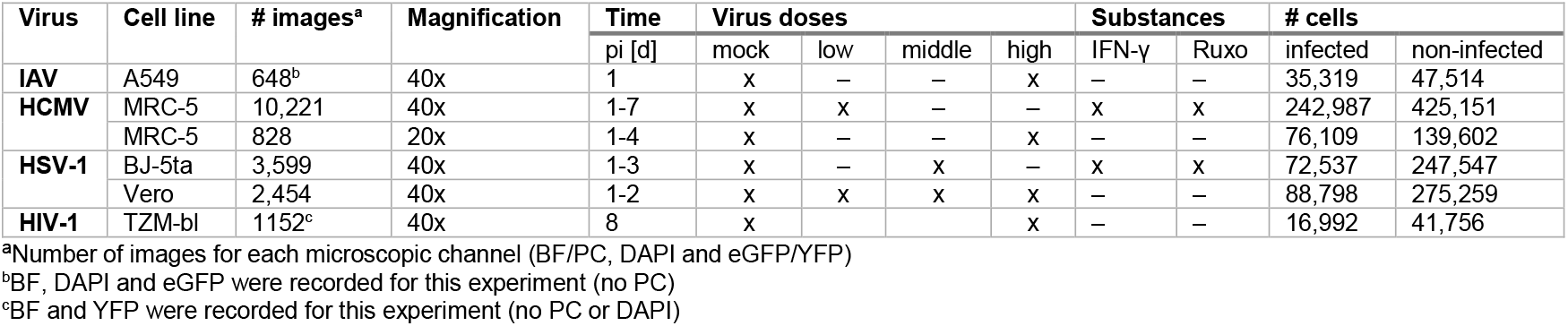
Image datasets of the examined viruses and cell lines.

Datasets were split into training, validation and test sets for the training of the DL model. The performance was evaluated for all virus and cell combinations using their respective test sets that were not used during the training process. For the first stream, tasked with predicting the signal of the fluorescent protein, the classification accuracy was calculated, along with the precision, sensitivity, specificity and negative predicted value (NPV). For this purpose, the eGFP/YFP intensity at the ground truth (GT) center based on the DAPI signal or BF cell position was determined using the original images, and the cells were classified as infected and non-infected accordingly (Figure 2A). The accuracy of the model demonstrates a strong performance, achieving values ranging from 83 % to 96 % for all viruses and cell lines. It is notable that the other metrics also achieve similar values for the corresponding combinations, indicating no substantial deficits in the detection of true positives (TP), false positives (FP), false negatives (FN) and true negatives (TN).

**Figure 2.**
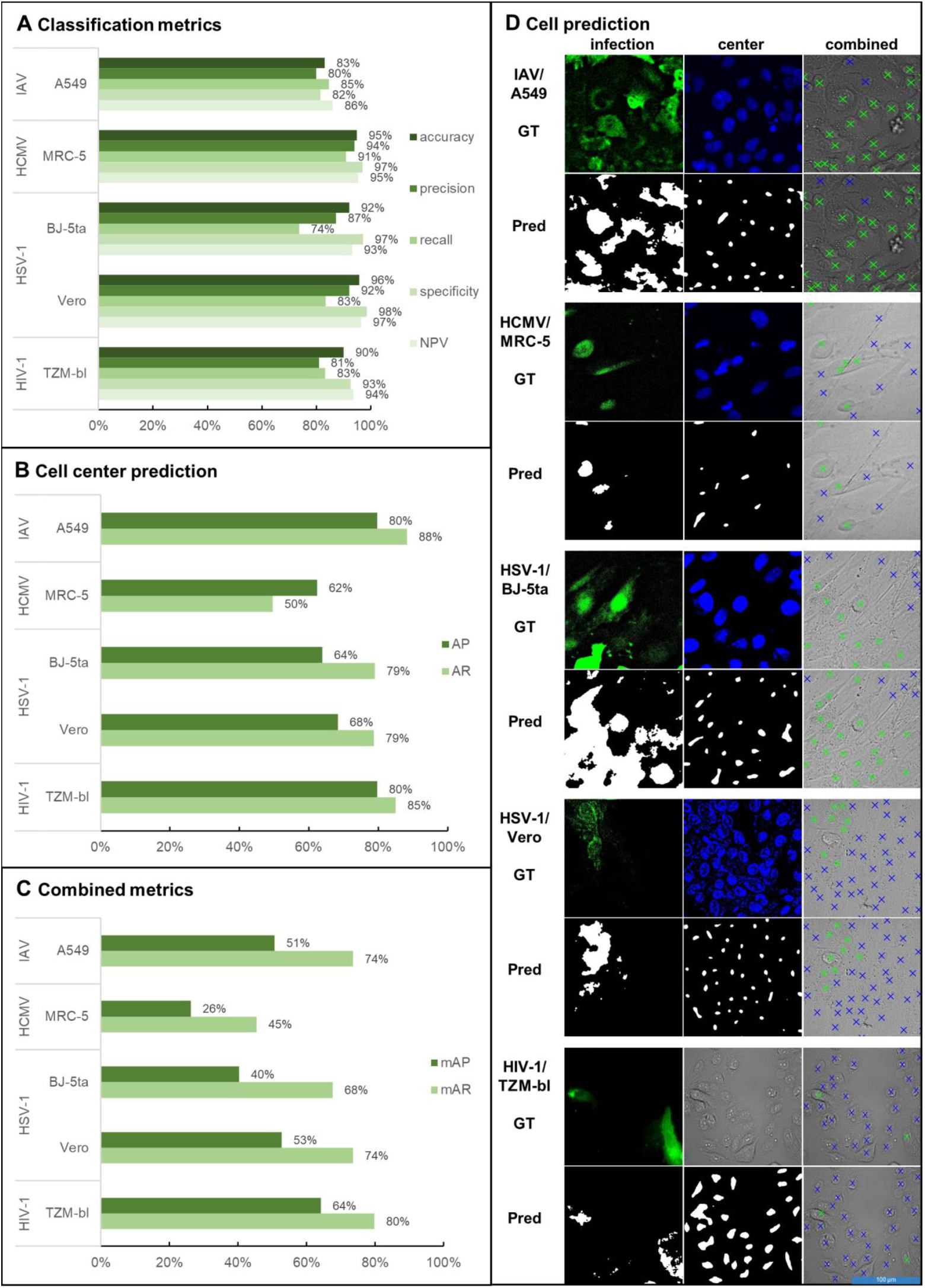
Classification metrics for infection status prediction, accuracy of center prediction, combined evaluation and visualization of GT and VAIruScope predictions. (A) Classification metrics (accuracy, precision, recall, specificity, NPV) were determined by comparing cells classified as infected (based on eGFP/YFP intensity at GT centers) with those classified based on the predicted signal for IAV, HCMV, HSV-1 and HIV-1 in the cell lines A549, MRC-5, BJ-5ta, Vero and TZM-bl. (B) AP and AR were used to determine the accuracy of cell center prediction by comparing predicted center locations with original DAPI/BF masks for IAV, HCMV, HSV-1 and HIV-1 in A549, MRC-5, BJ-5ta, Vero and TZM-bl cells. (C) mAP and mAR at an IoU of 10^-4^ were used to evaluate the accuracy of the overall model. Results are shown for IAV, HCMV, HSV-1 and HIV-1 in A549, MRC-5, BJ-5ta, Vero and TZM-bl cells. (D) Visualization of the eGFP/YFP intensity, the DAPI signal/BF image and GT classification, as well as the predicted infection signal, centers and classification (green crosses = infected, blue crosses = non-infected) for IAV, HCMV, HSV-1 and HIV-1 in A549, MRC-5, BJ-5ta, Vero and TZM-bl cells.

The accuracy of the second stream for center prediction was determined using the metrics average precision (AP) and average recall (AR). In order to accomplish this objective, a comparison was conducted to ascertain the presence of the predicted center locations within the original DAPI/BF masks (Figure 2B). The values demonstrate a high effectiveness, with APs ranging from 62 % to 80 % and ARs ranging from 50 % to 88 % for center prediction, as well as an adequate balance between the two values for the individual virus and cell line combinations.

In order to evaluate the performance of both streams in combination, the mean average precision (mAP) and mean average recall (mAR) were determined for existing intersection over union (IoU) for the predicted center and GT DAPI/BF masks, including their respective infection labels (Figure 2C). Average values for all combinations of 47 % for mAP and 67 % for mAR were achieved.

In Figure 2D, the eGFP/YFP and DAPI signals/BF images are displayed for an example of each virus and cell combination, with the resulting center coordinates and the assignment between infected (green crosses) and non-infected (blue crosses) cells overlaying the original BF/PC image. The predicted infection signal and centers are then shown below, along with the resulting classification and localization of the cells.

### Evaluation of infection kinetics, the effect of anti-viral and pro-viral drugs and virus dose

Next, we sought to test if VAIruScope can be utilized in experimental setups that are more reflective of common applications in virological and biomedical research. In order to investigate viral replication kinetics using VAIruScope, we analyzed cells at different time points, treatments and concentrations (Table 1). In Figure 3 the results for 1 to 7 dpi, anti-viral (IFN-γ) and pro-viral (Ruxo) drugs and virus doses (mock, low, middle, high) are shown. The percentages were compared with the GT values defined by eGFP signals.

**Figure 3.**
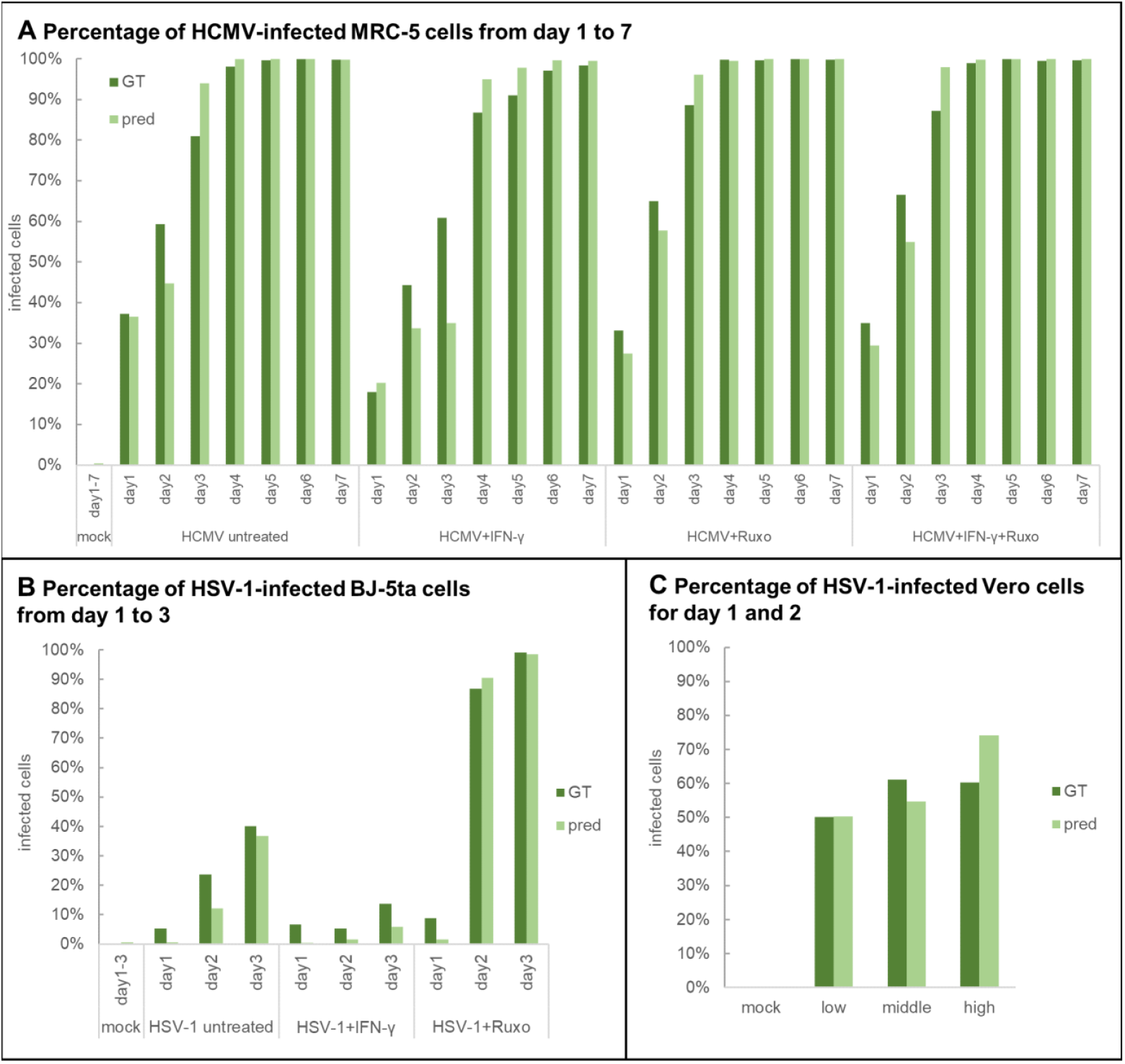
Percentage of HCMV- and HSV-1-infected cells for different time points, drugs and virus doses. (A) Percentage of HCMV-infected MRC-5 cells from day 1 to 7 and upon application of anti-viral (IFN-γ) and pro-viral (Ruxo) substances. Between 20 and 100 images were analyzed for each condition (time point and drug). The proportion for the GT classification and the prediction using VAIruScope is shown. (B) Percentage of HSV-1-infected BJ5-ta cells from day 1 to 3 and upon application of anti-viral (IFN-γ) and pro-viral (Ruxo) substances. About 30 images were analyzed for each condition (time point and drug). The proportion for the GT classification and the prediction using VAIruScope is shown. (C) Averaged percentages of HSV-1-infected Vero cells at different infection doses (mock, low, middle, high). Between 40 and 75 images were analyzed for each condition (virus dose). The proportion for the GT classification and the prediction according to VAIruScope is shown.

The proportion of cells being classified as infected by VAIruScope was consistent with the percentage based on the fluorescence signal for the majority of conditions, with an average deviation of 1 % to 8 % across all virus and cell line combinations. The prediction of non-infected cells in the mock condition demonstrated the largest accuracy ranging between 0.03 % and 0.6 % false positive classified cells.

The study also demonstrated that anti-viral and pro-viral effects can be measured. Accordingly, IFN-γ substantially reduced the number of cells infected with HCMV and HSV-1, whereas Ruxo caused a considerable increase. For HCMV/MRC-5, Figure 3A illustrates that, the infection rate is diminished in the presence of IFN-γ relative to that of untreated cells, resulting in the proportion of infected cells reaching saturation on day 5 to 6 instead of day 4. As expected, based on the JAK dependency of IFN-γ signaling, the combination of both substances reverted the anti-viral effect of IFN-γ. The results of Figure 3B demonstrate a consistent pattern of behavior for BJ5-ta and HSV-1, but with distinctly lower infection rates in comparison. Furthermore, the accuracy of the prediction of infected cells is less precise at early stages, with an average deviation of 4 % for all timepoints. The dose-dependency of HSV-1-infected Vero cells is shown in Figure 3C. Here, minimal deviations of 5 % on average from the GT were found.

### VAIruScope validation by correlative light and electron microscopy using an authentic HSV-1 isolate

In order to assess if VAIruScope can be applied successfully for unmodified wild-type (wt) viruses and without fluorescence staining, Vero cells were infected with an authentic wt-HSV-1 isolate, fixed and readily subjected to microscopy imaging. Four different conditions (mock, low, middle and high infection doses) were examined. Figure 4A shows plots of the percentage of predicted infected cells for the different doses of wt-HSV-1. For the mock condition, less than 1 % were false positive for the untreated cells. Dose-dependency of wt-HSV-1 infection is depicted in Figure 4B.

**Figure 4.**
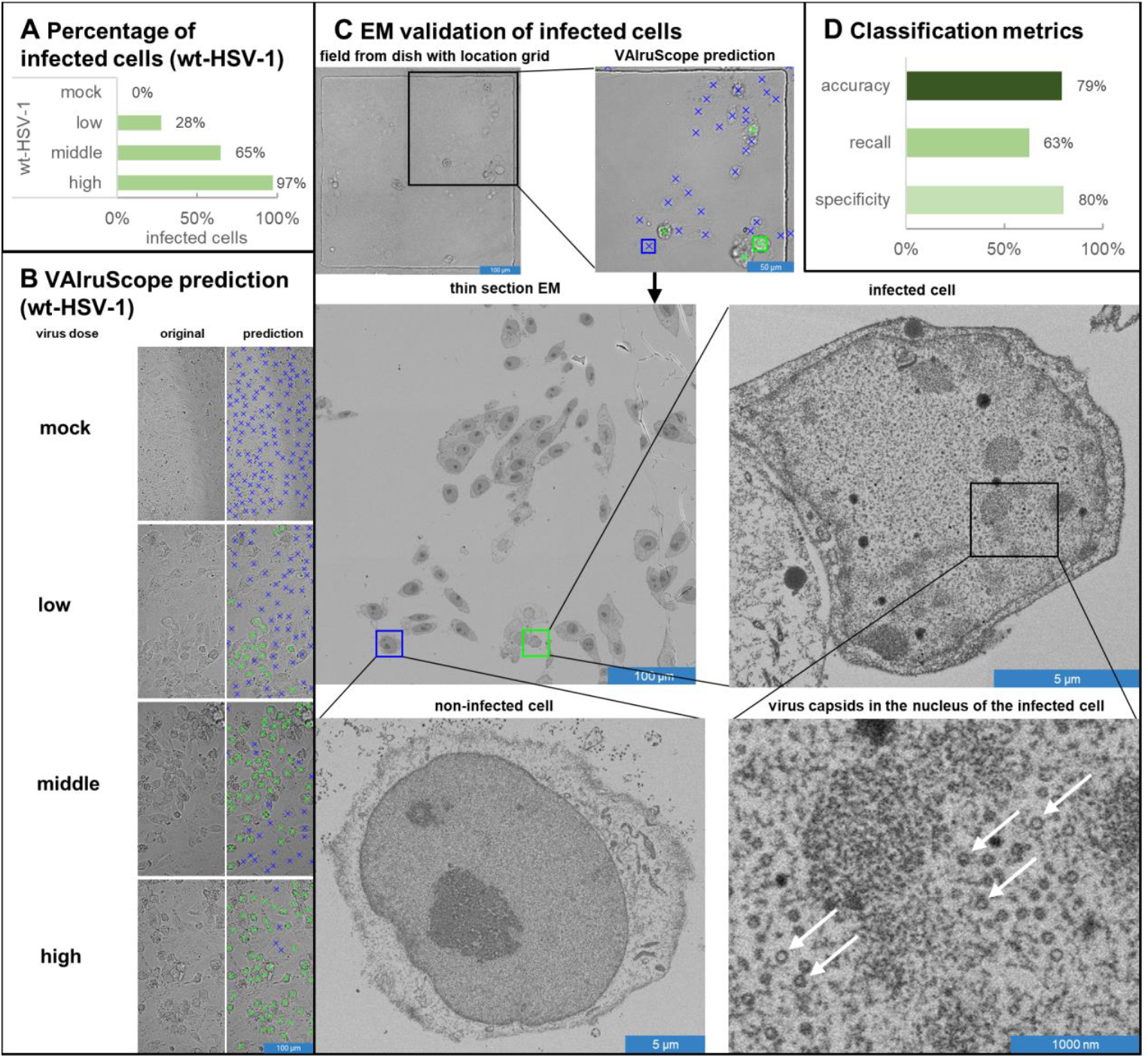
wt-HSV-1 infection and EM evaluation. (A) Percentage of wt-HSV-1-infected Vero cells at different conditions (mock, low, middle, high infection doses: see details in the method section) according to VAIruScope predictions. 105 images were analyzed for each condition. (B) Visualization of the prediction of wt-HSV-1-infected (green crosses) and non-infected Vero cells (blue crosses) according to VAIruscope for different conditions (mock, low, middle, high infection doses). (C) The wt-HSV-1-infected Vero cells in a dish with a labelled grid pattern. A distinct mesh of the grid is shown by BF microscopy, and for a subregion the predictions by VAIruScope are indicated in the overlay. The subregion is shown by thin section EM (one section of the section series through the entire volume of the cells is shown). A non-infected and wt-HSV-1-infected Vero cell are shown at higher magnification. Infection of the cell is expert-rated based on the presence of the assembling virus capsids in the cell nucleus (white arrows). (D) Accuracy, recall and specificity of the classification using VAIruScope compared to EM results. Two grid cells with a total of 158 cells were manually examined.

We then used correlative light and electron microscopy (CLEM) with thin sections to evaluate the VAIruScope predictions. To this end, cells were cultured and infected in a dish with a labelled grid pattern. Individual cells were imaged and predicted based on the BF channel using VAIruScope. To confirm the assignment, cells were inspected regarding the presence of HSV-1 particles in their nuclei using CLEM. Figure 4C shows a distinct mesh of the grid by BF microscopy, with the prediction obtained by VAIruScope, and expert-rated based on the serial thin section EM. An infected cell with virus capsids in the cell nucleus (ring-shaped particles, marked with white arrows) and a non-infected cell are shown at higher magnification as examples. A total of 158 cells were evaluated using both methods (Figure 4D). An accuracy of 79 %, a slightly lower recall of 63 % and a specificity of 80 % were achieved, which is comparable to the values determined in Figure 2A for HSV-1-infected Vero cells.

## Discussion

In this study, we introduce VAIruScope, a label-free tool for the detection of infected cells based on BF/PC microscopy that is suitable for DNA-, RNA- and retroviruses and opening up the possibility for the rapid and cheap investigation of either infection dynamics, the effects of drugs appropriate for wt and even emerging viruses. Our approach requires no fluorescent staining, works with standard light microscopy and reliably detects infected cells just one dpi, allowing for the investigation of early timepoints and low viral doses. In addition, a single model can be used across various virus/cell combinations.

### Deep learning label-free detection of infected cells

We developed a versatile DL pipeline, VAIruScope, for the label-free identification of virus infection at the cellular level with high accuracy.

As indicated by various evaluation metrics for infection classification, center prediction, and combined analysis (Figure 2A-C), the model delivers reliable results for four different viruses in five different cell lines. There were two major challenges in the development of the model. Firstly, the infection-related eGFP/YFP signal depends on various factors including the exact time post infection for each cell, the specific virus-encoded reporter gene, the protein distribution in the cell and is overshadowed by noise of the microscopic setup. Given these variations, the exact definition of an infected cell based on the fluorescence signal remains ambiguous. In order to include a variety of cells during different stages of the infection cycle, we choose a global threshold per virus and cell combination independent of dpi, concentrations, or treatment. In future work, the investigation whether different stages of infection could be separated or not, should be examined, but would require more controlled and synchronized infection conditions. The second challenge lies in the high variability of the different cell lines and the imbalance of infected cells per image. Setting up the model to include all cell lines and viruses in a joint training set greatly benefited the overall training stability. The identification of cell centers was particularly improved by training the model using cells from the different cell lines, which allowed it to learn shared morphological features. However, the center predictions show some variability with MRC-5 having the lowest accuracy, potentially due to the rather irregular morphology of this cell line.

As shown in Figure 3A-C, VAIruScope accurately reflects the dpi, the effect of drugs on viral infectivity and the influence of different virus doses. As expected, the kinetics for HCMV/MRC-5 and HSV-1/BJ-5ta showed a dynamic increase of infected cells in relationship to the initial infection dose. However, for the HSV-1/BJ-5ta combination, the number of infected cells is slightly underestimated for most time points, also reflected in the lower recall value for that virus/cell combination. Nevertheless, the overall offset remains consistent across all time points, clearly reflecting the trend in infection dynamics and drug-induced alterations.

In summary, VAIruScope demonstrates a solid capacity to accurately identify individual virus-infected cells from only BF/PC microscopic images, thus overcoming the limitations associated with conventional methods that necessitated specific reporter viruses and/or staining procedures.

### Comparison with existing approaches

Previous approaches have either been limited to examining the level of individual cell culture wells [20–26] or require specialized microscopic or staining techniques [18, 27, 34], limiting their applicability. Therefore, viral replication or infection dynamics could only be evaluated by capturing a population of cells. Furthermore, specialized microscopic devices are expensive and cumbersome to use and staining techniques often rely on fixation and therefore exclude investigations of infection kinetics.

It has been demonstrated that DL image classification approaches (e. g. custom convolutional neural networks [20], ResNet [21, 35]), which examine the CPE at whole culture wells capturing hundreds of cells simultaneously, can achieve classification accuracies of over 90 % (e. g. Wang, et al. [20] for IAV, Werner, et al. [21] for SARS-CoV-2) after a period of one to three dpi and high multiplicity of infection (MOI). Utilizing a comparable methodology, Dodkins, et al. [22] successfully classified six distinct viruses early after infection (4-24 hours post infection (hpi)) with the MobileNet model [36]. Its accuracy ranges from 74 % to 97 % and reaches a maximum of 80 % for certain viruses (HIV-1, coronavirus-229E, vaccinia virus). Two research groups developed multitask classifiers (ResNet [23, 35], EfficientNet-B0 [24, 37]) for six [23] to eight [24] viruses with accuracies of 80 % to 98 %. The training was conducted on images of cell cultures with advanced dpi, relying on a well-developed CPE. A study on the detection of vesicular stomatitis virus (VSV) demonstrated over 99 % accuracy by analyzing the cell confluency as an indirect, correlated measure of infection [25]. As a further development of this research, researchers found that cell rounding is directly proportional to the virus dose for three different viruses (VSV, Newcastle disease virus, parapoxvirus ovis), allowing for precise titer determination [26]. Both methods necessitate the utilization of an advanced Cytation5 microplate imager or proprietary software [25, 26]. Although DL has improved infection detection in cell culture, current approaches still face key limitations, including virus-specific models, dependence on specific (often high-titer or late) infection conditions, costly specialized imaging and proprietary software, and the absence of single-cell resolution, which together hinder detailed analysis of infection dynamics and drug effects.

Unlike whole-well-based methods, some approaches analyze individual cell images for CPE: Yakimovich, et al. [27] used digital holo-tomography microscopy and refractive index gradient measurements to detect CPEs of vaccinia virus, HSV-1 and rhinovirus, while another study employed Raman spectroscopy to reliably identify HSV-1, HSV-2 and varicella zoster virus-infected cells [34], both methods rely on specialized equipment or continuous time-series acquisition. The findings of Andriasyan, et al. [18] are most comparable to our methodology. In that study, the authors examined HSV-1- and adenovirus (AdV)-infected A549 cultures at the single-cell level using the nuclear fluorescence signal. The method could be used to detect the morphological changes of the nuclei associated with cell infections. Employing the ResNet architecture [35], a cell-specific classification accuracy of 94 % for HSV-1 was achieved at two and three dpi. This accuracy is comparable to our results of HSV-1-infected BJ-5ta and Vero cells with an accuracy of 92 % (one to three dpi) and 96 % (one to two dpi), respectively. However, despite having a similar detection accuracy, VAIruScope only uses the BF/PC image as input. This omits fluorescence staining, thereby reducing costs and manual work, as well as the need for intervention with the system. This broadens its general applicability.

### Potential applications for VAIruScope

VAIruScope offers a promising foundation for label-free detection of virus-infected cells, with a wide range of applications in various fields. It is a valuable tool that facilitates basic research in the domain of infection dynamics and network effects. VAIruScope can be used to study viral pathogenesis and replication during label-free live-cell imaging and to evaluate the effect of anti-viral drugs and vaccine-induced antibodies. Following the identification of infected and non-infected cells, specific molecular biological methods (e. g. PCR, next-generation-sequencing) can be employed to identify factors that render cells more or less permissive to viral infections that can lead to the detection of relevant host factors. As some commonly used laboratory strains harbor adaptations to cell cultures [38–42], reproducing key findings with clinical virus isolates and/or the investigation of authentic pairs of viruses with mediators from the genuine host (e. g., virus and immune serum pairs) is becoming increasingly important. Additionally, it is suitable for the investigation of wt viruses, potential novel strains or even newly emerging viruses as it requires no genetic modification of the virus, nor does it depend on specific antibodies or primer sequences. We investigated wt-HSV-1-infected Vero cells with VAIruScope showing virus concentration-dependent results (Figure 4A and B). In addition, we compared the predictions with an EM-based validation, showing a strong correlation (Figure 4C and D).

### Limitations of the study

The developed DL framework VAIruScope has been demonstrated to possess the capability to accurately identify virus infections on microscopic BF/PC images at the single-cell level. The final method is label-free, but most of the training data was conducted using DAPI-stained cell nuclei and genetically modified viruses to determine the infection status. Using this approach IAV-, HCMV-HSV-1- and HIV-1-infected cells can be detected and wt-HSV-1 infection was validated using EM.

It has been demonstrated that the developed methodological approach is also well suited for BF and PC microscopy images and different magnifications. However, a variation in the imaging setup, cell line or virus strain might reduce the detection accuracy. Further fine-tuning of the model using new training data for a modified configuration could lead to a corresponding adaptation.

The results for HSV-1 show that the accuracy of the classification is not uniform for different cells (Figure 2A). These findings suggest that CPE may be stronger or more variable between different cell lines infected with the same virus. Consequently, additional training of VAIruScope would be required to optimize its use in new combinations. Nevertheless, in such scenarios, the employment of transfer learning techniques [43] has the potential to rely on smaller datasets comprising meaningful examples for novel use cases. Since VAIruScope has already been shown to achieve reliable results for four different viruses and five cell lines, adapting it for new combinations is likely to be straightforward. Nevertheless, the transferability and generalizability of VAIruScope would need to be investigated. It is conceivable that the DL-based approach could detect morphological changes that are barely visible to the human eye in light microscope images.

Our method demonstrated that the developed techniques are effective for chemically fixed cells. However, to analyze the infection dynamics or replication mechanisms of viruses in detail, it is necessary to examine the infection in live-cell-imaging.

## Methods

### Cell culture

A549 (DSMZ, ACC 107), MRC-5 (ATCC. CCL-171), BJ-5ta (ATCC. CRL-4001), Vero (ATCC. CCL-81) and TZM-bl (RRID: CVCL_B478) cells were grown in high glucose Dulbecco’s minimal essential medium (DMEM [Gibco 41966-029]), supplemented with 10 % (v/v) FCS. For MRC-5, BJ-5ta and Vero cells medium was supplemented with 100 μg/ml streptomycin/100 U/ml penicillin (Gibco). Cells were kept at 37 °C in an atmosphere of 5 % CO_2_. A549, MRC-5, BJ-5ta and Vero cells were regularly screened for mycoplasma through PCR (Minverva, VenorGeM) and TZM-bls with the InvivoGene MycoStrip Kit. All cell lines were only used if they were free of mycoplasma.

### Viruses, infection and treatment

A549 cells were infected with influenza A virus PR8-NS1-GFP at MOI1 (high virus dose) in serum-free DMEM supplemented with 0.2 % BSA (Sigma). Cells were infected for 1 h before the medium was changed to serum-free DMEM supplemented with 0.2 % BSA (Sigma). MRC-5 were infected with virus doses 0.1 (low) and 1 (high) PFU/cell of HCMV TB40-eGFP (described in PMID: 24337170) applying centrifugal enhancement (900 g, 30 min at RT). Vero cells were infected with the virus doses 0.01 (low), 0.1 (middle) and 1 (high) PFU/cell of HSV-1-ΔgE-GFP (generated and described in PMID: 12857917, provided by the laboratory of Prof. David C. Johnson, Oregon Health & Science University, USA) or an authentic HSV-1 isolate (0.05 (low), 0.1 (middle), and 1 (high) PFU/cell, PMID: 21264213). For treatment experiments, BJ-5ta cells were pre-incubated for 15 min with either Ruxolitinib ([4 µM]; Cell Guidance Systems) or human IFN-γ ([100 IU/ml], PBL) and then infected with HSV-1-ΔgE-GFP (0.1 (middle) PFU/cell). TZM-bl cells (5 × 10^4^) were infected with a modified version of recombinant HIV-1 construct (HIV_DFIII_), generated and described in Sperber et al. (PMID: 37910505). Here, a membrane-associated YFP replaced the GFP of the original construct, so that virus infected cells with active HIV-1 replication could be identified based on their YFP signals. Prior to infection, the virus was titrated, and infection efficiency was determined via flow cytometry. In all experiments, a low virus titer was used to maintain a multiplicity of infection (MOI) below 1. Infections were performed via spinoculation (997 g, 2 h at 37°C). At 24 h post-infection the medium was changed to DMEM supplemented with 10% FBS and 100 μg/ml streptomycin and 100 U/ml penicillin.

### Brightfield and fluorescence microscopy

To generate the images for classification, mock or IAV/HCMV/HSV-1-infected cells were fixed at the indicated time points with 4 % (w/v) paraformaldehyde/PBS at RT for 20 min. Cells were washed twice with PBS and permeabilized with 0.2 % (v/v) Triton-X-100/PBS for 15 min at RT. Cells were washed again with PBS and stained with 4′6-diamidino-2-phenylindole (DAPI; Sigma) for 3 min at room temperature (RT). Cells were washed twice in PBS and stored in PBS at 4 °C until imaging. TZM-bl cells were washed once with PBS and subsequently fixed 8 dpi using 4% (w/v) paraformaldehyde in PBS for 15 minutes at RT. After fixation, cells were washed with PBS and stored in PBS at 4 °C until imaging.

Microscopic images were acquired with Leica Thunder 3D Cell Culture using the LASX 3.7.6 software with a magnification of 400 (10x Ocular; 40x air objective; NA 0.6) or a magnification of 200 (10x Ocular; 20x air objective; NA 0.4) and LAS X software 3.7.6. with automatic background correction (ICC).

### Correlative light and thin section electron microscopy (CLEM)

Vero cells were plated in a cell culture dish with a grid pattern in the plastic bottom (µ-Dish, 35 mm, Grid-500, ibidi) and infected with a clinical HSV-1 isolate (1 PFU/cell). At 24 hpi, cells were fixed, permeabilized and stained with DAPI as described above (Brightfield and fluorescence microscopy).

After BF microscopy, cells were additionally fixed with 2.5 % glutaraldehyde in 0.05 M Hepes buffer for several hours. Cells were post-fixed and contrasted with 4 % osmium tetroxide (60 min), 0.2 % tannic acid (60 min), 2 % uranyl acetate (60 min) and 0.67 % lead aspartate (30 min) at 60 °C. After dehydration in ethanol, cells were infiltrated with epon resin using acetone as intermedium. The infiltrated cells were overlaid with a thin layer of epon (up to the inner rim of the dish) and covered with a sheath of Aclar plastic film. Finally, the epon was polymerized at 60 °C for 2 days. Regions of interest were identified by their grid position and extracted from the polymerized layer by using a hot scalpel. After mounting the extracted part to an aluminum stub, the bottom was removed with a histo-cryo diamond knife (Diatome) at the ultramicrotome (UC7, Leica Microsystems) and serial sections (90-120 nm thickness) were taken parallel to the bottom using a standard diamond knife for ultrathin sectioning (Diatome). Sections were collected on silicon wafers, glued on an aluminium stub and coated with carbon. Imaging was done with a scanning electron microscope (Teneo VS, Thermo Fisher) and the T1 detector at 2 kV using the MAPS software (Version 2.5, Thermo Fisher). Cells were identified in overview images and inspected at every section level for the presence of virus structures by two trained specialists.

### Dataset preparation

The size of the datasets is subject to variation due to the number of microscopic images, the time period examined, and virus doses used for infections, thereby enabling the investigation of a variety of different conditions for the detection of infected cells (Table 1). Either BF or PC images were selected for analysis depending on their quality or availability (BF for IAV/A549, HCMV/MRC-5 20x, HSV-1/BJ-5ta and Vero, HIV-1/TZM-bl, PC for HCMV/MRC-5 40x). The microscopic BF/PC and fluorescence images of the individual experiments were compressed into a stack with two to three image channels for further analysis. All images were square in shape and resized to a standard dimension of 1024×1024 pixels (px) with 16-bit pixel depth. The image datasets were split into training, validation and test sets (80:10:10) for each virus/cell line combination.

Either the DAPI channel of the image data (IAV, HCMV, HSV-1) or the BF channel (HIV-1) was utilized for the purpose of detecting the cell centers, employing the Cellpose nuclei (nucleitorch_0) [44] or the Cellpose-SAM model [45]. It is important to note that nuclear staining was not performed for the TZM-bl/HIV-1 combination in order to avoid any possible interference with the infection signals. The DAPI and cell masks that are produced subsequently serve the process of calculating their center. The center coordinates based on image moments are converted into Gaussian peaks, whose intensity correlates with the size of the nuclei or cell. The resulting masks are used for model training.

In order to normalize the fluorescence values of the virus-related proteins, a background subtraction was first performed. To accomplish this, the average intensities in the mock images were determined and subsequently subtracted from all images of the corresponding experiment. In addition, the eGFP/YFP signal values were further normalized based on the maximum measured value for each experiment. Afterwards, the mean intensity of the mock cells was ascertained within a 5×5 px box, based on the calculated cell centers. An intensity threshold for infected status definition was selected for each experiment, such that 99 % of the mock centers (except for IAV/A549 and HIV-1/TZM-bl, for which a value of 90 % and an intensity greater than 1 was applied) exhibit a lower average signal in the channel of the fluorescent protein. The normalized eGFP/YFP images were converted into binary masks based on the calculated intensity threshold from the mock condition. For HSV-1 and HCMV, the cell center can be used as a key feature for virus identification, as it is assumed that the accumulation of the eGFP is most intense in the nucleus [46]. In contrast, for IAV it is expected that eGFP will be primarily expressed in the cytosol [47]. To ensure that the cell center could still be used as the basis for determining cell infection, DAPI masks were added to binary eGFP masks wherever they overlapped with infection areas. Nevertheless, eGFP/YFP images from the mock condition with intensities above the threshold were considered negative (and converted to all-zero masks) for model training and further analysis.

### DL algorithms and training

All implementations were carried out in Python using the PyTorch framework [48] and the Hugging Face Transformers library [49]. The model employed for differentiation between infected and non-infected cells and cell localization is a modification of the UPerNet architecture [50] using a ImageNet [51] pre-trained Swin-L transformer [52] backbone. Extracted hierarchical feature maps from the input images were subsequently processed by the UPerNet decoder head to produce intermediate outputs for both processing streams.

Following the image decoder, the network splits into the two streams. The first stream was designed to differentiate between infected and non-infected cells, using binary masks from the eGFP/YFP channel. The second stream of the model was trained to predict cell centers, using masks derived from the DAPI/BF channel. In each stream, the original input images are processed through two sets of U-Net-style [53] convolutional blocks, each comprising sequential convolutional layers with batch normalization and ReLU activations, followed by pooling and upsampling operations to extract multi-scale features. The upsampled decoder outputs for each stream are concatenated with the corresponding down sampled features, and the resulting fused representations are further refined through the remained convolutional blocks. The final convolutional layers of each stream produce two separate output maps: one for classification between infected and non-infected cells and one for cell localization.

Each stream is optimized using a distinct loss function, namely, binary cross-entropy with logits (BCE) for the first stream and mean squared error (MSE) for the second. Training images were augmented with random horizontal and vertical flips and rotations to improve generalization. In total, 7,816 BF and 9,862 PC images of the different cell lines (incl. virus infections) with corresponding labels for both streams were used for training, and 1,887 BF/PC images were reserved for testing.

The network was trained for 25 epochs with a total batch size of 16, distributed across four H100 GPUs. The AdamW optimizer [54] was used in conjunction with a linear learning rate scheduler, incorporating a warmup period of two epochs. Training and validation loss were defined as a combination of the BCE loss from the first stream, weighted by 2 and the MSE loss from the second stream, weighted by 0.05:

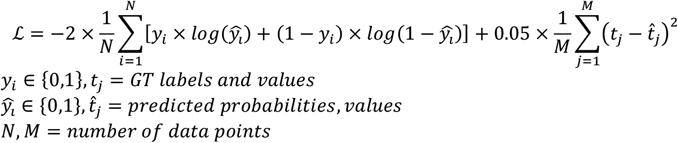

The model achieving the lowest validation loss was selected for further analysis.

This design allows the model to leverage global contextual representations from the Swin-L backbone while simultaneously refining local, high-resolution details through the U-Net based modules, effectively combining coarse predictions with fine-grained features to improve overall performance.

The predictions of both streams were evaluated separately and in combination on the test sets. After entering a BF/PC image, the predicted mask of the first stream was evaluated using the GT centers. The coordinates were employed to measure the infection signal in a 5×5 px area around the center. If the value exceeded a mean of 0.5, the cell was designated as infected (for the CLEM experiment, a value of 0.9 was used to focus on the prediction of highly infected cells). The cells classified by this method were compared with the cells differentiated by the original label (consisting of eGFP/YFP signal and overlapping DAPI/BF location), and the following metrics were determined:

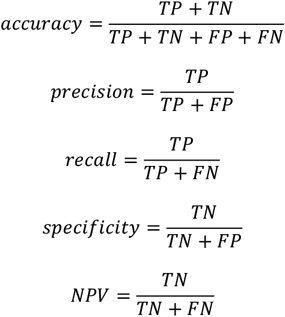

From the predicted mask of the second stream, local maxima were identified and treated as individual cell centers, whose coordinates were subsequently calculated. The Cellpose metrics code [44] was used to calculate the AP and AR at an IoU of 10^-4,^ rating any overlap of the predicted center point with the corresponding GT DAPI/BF mask as correctly identified.

Finally, the results of both streams were merged after training – the predicted infection masks from the first stream were used in combination with the calculated centers from the predicted masks from the second stream to localize and distinguish between infected and non-infected cells. A polygon with an area of 100 px^2^ was delineated around the center coordinates, and was saved together with the predicted label (infected/non-infected). A comparison was made with the GT DAPI/BF masks and their corresponding infection labels. The performance of VAIruScope was determined using the metrics mAP and mAR at an IoU of 10^-4^ and a maximum of 1000 detections aided by the common objects in context (COCO) evaluation code [55].

## Declarations

### Resource availability

Code and data are available upon reasonable request from the corresponding author.

### Author Contributions

Conceptualization, Ch.S., M.T., R.S. and N.K.; methodology, J.P., Ch.S., M.T., R.S. and N.K.; software, J.P., S.A. and A. L.; validation, A.K. and M.L.; investigation, J.P., Co.S., E.M., A.L., A.K. and M.L.; resources, Co.S., E.M., T.T., B. G., A.L. and VTK.LT.; writing—original draft, J.P., Co.S. and N.K.; writing—review & editing, Ch.S., M.T., R.S. and N.K.; supervision, Ch.S., M.T., R.S. and N.K.; funding acquisition, N.K.;

## Acknowledgements

We would like to express our profound gratitude to Joana Grah for helpful discussions related to the model architecture. We also thank Gudrun Holland for resin embedding of the cells for CLEM.

